# Comparative Analysis of the Techniques for the Determination of Binding Affinity between a Small Molecule Inhibitor and a Protein Target

**DOI:** 10.1101/2024.05.16.594462

**Authors:** Yalan Luo, Yijun Chen

## Abstract

The binding affinity constant (*K*_*D*_) between a small molecule inhibitor and a protein target is pivotal parameter for target identification and early drug discovery. Despite the extensive applications of three major techniques, namely SPR, ITC and MST, the *K*_*D*_ values for a specifically defined binding measured by these techniques are drastically different, which poses a remarkable difficulty to deal with these ambiguous data. Here, we report the evaluation of the accuracy of *K*_*D*_ values from SPR, ITC and MST compared with enzyme kinetics. To enable an objective comparison, we theoretically proved that enzyme competitive inhibition constant (*K*_*i*_) could directly reflect the binding affinity. Using purine nucleoside phosphorylase, its substrate inosine and competitive inhibitor immucillin-H, we determined respective *K*_*D*_ and *K*_*i*_ values to make a direct comparison. Moreover, we found that the *K*_*D*_ value measured by SPR is more relevant to its *K*_*i*_ value. This study highlights the urgent need on the development of new technologies for the determination of binding affinity between small molecule inhibitors and protein targets.

## 1. Introduction

As commonly known, binding affinity represents the strength of ligand-protein interaction, which quantifies the extent of protein occupancy. Generally, affinity calculation is based on following three methods: a. Δ*G*_*binding*_ *= RTln(K*_*D*_), which relies on thermodynamics [1]; b.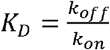, which depends on kinetics [2]; and c.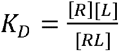, which is based on equilibrium dynamics [3]. These formulas are calculation basis of different detection methods for affinity, but they all principally rely on the binding affinity between dissociated components [2, 4]. When the interaction of a small molecule inhibitor and its target protein is concerned in the case of early drug discovery, it is preferable to accurately measure inhibitor-target affinity because it is an indicator to associate with the efficacy of inhibitors and their off-target effects. Consequently, as an important parameter, binding affinity constant (*K*_*D*_) has been a valuable *in vitro* measure for various preclinical evaluation, including the selection of lead compounds, the optimization of leads, and the predictions of *in vivo* potency, pharmacokinetic properties and safely profiles [5]. Therefore, it is of great important to obtain accurate *K*_*D*_ values of the inhibitors.

Despite the complexity of biochemical reactions in all living processes, every type of physical and chemical transformations results in a series of changes in dynamics and thermodynamics. As a result, the emerging technologies for the determination of binding affinity between different molecules are literally designed to measure these changes [6]. Currently, there are three major techniques for this purpose, namely Microscale Thermophoresis (MST), Isothermal Titration Calorimetry (ITC) and Surface Plasmon Resonance (SPR). MST determines binding affinity through the variation of thermophoresis of a fluorescent labeled molecule, and such a variation is usually caused by the changes of molecular properties, including size, charge, hydration state, and/or structural/conformational changes of the proteins, under constant buffering conditions upon the binding with small molecule inhibitors [7]. ITC focuses on the thermodynamics of molecular interaction where the contacts between two molecules produce exothermic or endothermic reaction to generate the information on association equilibrium constant, enthalpy change and stoichiometry from the binding [8]. The principle of SPR utilizes optical phenomena for detecting the association and dissociation of analytes on a biosensor chip by real-time monitoring [9]

In recent years, the binding affinity values and their measurement techniques have greatly contributed to accelerating the process of target identification and early drug discovery. Unfortunately, as a conventional wisdom and common practice, the choice of using which technique largely depends upon the availability of instrumentation rather than data reliability, resulting in unprecedented uncertainty on the reliability and/or reproducibility of *K*_*D*_ values for a pair of ligand-protein across different techniques [10]. Especially, for the interaction between high weight proteins and small chemical compounds (typically molecular weight < 500), the significant difference on molecular weight of the analytes leads to inconspicuous signal from the interaction, causing an exceptional discrepancy. For example, ABT -737 is one of the inhibitors of BCL-2, and its binding affinity to BCL-2 was determined by SPR and ITC to be 0. 6 nM and 20.5 nM, respectively [11, 12]. The affinity of MCL-1 inhibitor A-1210477 was reported as ∼740 nM by MST [13], whereas another study showed 3.5 nM by SPR [14], an over 200-fold difference. More intriguingly, using the same technique for the same interaction produces even larger difference, such as the case between PDL-1 and BMS-202 measured by MST to be 1.1 μM and 37 nM in two different investigations [15, 16]. Therefore, these obviously discrepant data have generated a serious question on the reliability of reported *K*_*D*_ values and their associated techniques.

When enzyme kinetics is concerned, the inhibition constant (*K*_*i*_), particularly in the case of competitive inhibition, is extremely similar to *K*_*D*_ value to represent the binding affinity. *K*_*i*_ is a thermodynamics parameter, an important parameter and gold standard for the efficiency of inhibitors, and the true affinity between an inhibitor and an enzyme [17]. Previous studies have also verified that binding affinity and enzyme inhibition are closely correlated in thermodynamics, where *K*_*i*_ values strongly correlate with *K*_*D*_ values. Meanwhile, competitive *K*_*i*_ is quite close to *K*_*D*_ when the structures of substrate and inhibitor are remarkably similar [18]. Thus, we speculate that competitive *K*_*i*_ can serve as the reference to estimate *K*_*D*_ under consistent and reproducible conditions. In this study, we demonstrated the close correlation between competitive *K*_*i*_ and *K*_*D*_ by a comparative analysis, which could facilitate the development of more reliable technologies for *K*_*D*_ determination.

## 2. Methods and Materials

### 2.1 Expression and purification of human purine nucleoside phosphorylase (hPNP)

BL 21 (DE3) cells were transformed with pET28a (+)-hPNP plasmid and grown in 20 ml Luria-Bertani medium with kanamycin (50 μg/ml) overnight, then 2 ml culture was inoculated to 200 ml fresh LB medium. The diluted cultures were grown at 37°C to approximately *OD*_600_*=* 0.6 for the induction with IPTG (250 μM final concentration). After 20 h of additional growth at 20°C, cells (∼1 g) were harvested by configuration at 4 □.

Approximately 12 g cells were suspended in 50 mM potassium phosphate buffer (pH 8.0) containing 0.1 mM PMSF. After ultrasonication and centrifugation at 4□, the supernatant was incubated with Ni-NTA affinity resin for 1 h at 4 □. hPNP was eluted with 14 mL 50 mM potassium phosphate buffer (pH 8.0) containing 300 mM imidazole. Approximately 80 mg protein purified via Ni-affinity chromatography was loading on Q-High performance anion exchange chromatography column for further purification, in which the column was equilibrated with 50 mM HEPES (pH 8.0), and then hPNP was eluted with a linear gradient of 0-0.25 M NaCl. The purity of hPNP was examined by 12 % SDS-PAGE with Coomassie blue staining.

### 2.2 Enzyme assay and determination of competitive inhibition

The increase of uric acid was determined by SpectraMax iD5 instrument in a microtiter 96-well plate. The reaction mixture contained 1.4 nM hPNP, 25 μM to 5mM inosine as substrate and 0 -5 nM immucillin-H (ImmH) in the wells at 25 □. Michaelis-Menten equation was employed to obtain *K*_*m*_ values, and the equation 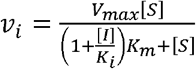 was used to generate *K*_*i*_ *values*. The data were analyzed by Graph Pad Prism9.

### 2.3 Binding affinity assays

SPR experiments were performed with a Biacore T200 instrument in 50 mM potassium phosphate buffer (pH 7.4) at 25 □. Purified hPNP was diluted to 20 μg/ml in 10mM sodium acetate solution (pH 5.5) and immobilized on the sensor chip to reach a final immobilization level of 7000 RU. ImmH was diluted to 1.2 - 925 nM with flow rate of 30 μl/min and contact time of 120 s in each injection cycle.

ITC experiments were performed with a PEAQ-ITC instrument in 50 mM potassium phosphate buffer (pH 7.4) at 25 □. The starting concentrations of hPNP and ImmH were 5 μM and 150 μM. One Sit of Sites model was used with a setting of Reference Power, FeedBack, Stir Speed, Initial Delay by default. In addition, injections and the volume of each injection and spacing time were 15, 2.5 μl and 150 s during each experiment.

For MST determinations, 200 nM hPNP was incubated with 100 nM fluorescent tags for 30 min in His-Tag Labeling Kit RED-tris-NTA 2nd generation to reach enough raw fluorescent counts (> 200). The concentration of ImmH was varied in a dilution series starting at 1 mM, and then mixed with fluorescent-labeled hPNP in 50 mM potassium phosphate buffer (pH 7.4). After loading the samples into standard capillaries, scanning was performed under 40 % MST power and 40% LED power at 25□.

## 3. Results

### 3.1 The relationship between enzyme competitive inhibition and biding affinity

Different types of reversible inhibitions in enzyme kinetics exhibit various relationships between 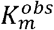 and *K*_*i*_ (Scheme 1a) [19]. Among the inhibitions, competitive inhibition provides the simplest and an intuitive linear relationship:

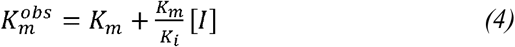

In competitive inhibition, a substrate competes with an inhibitor to bind the active site of the enzyme. There are three equilibriums in competitive inhibition (Scheme 1b):

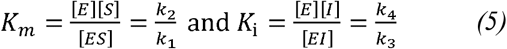

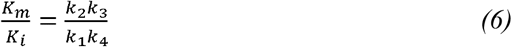

Apparent Michaelis constant 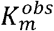 contains two equilibriums and can be calculated in accordance with Eqs. (1) and (3). Therefore, apparent inhibition constant 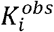 could be derived from Eq. (2) and reverse equilibrium of Eq. (3). Subsequent rearrangement generates Eqs. (7) and (8):

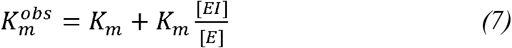

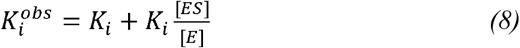

Apparently, the *K*_*i*_ value from theoretical calculation represents the initial binding state of an enzyme and an inhibitor, which is the true affinity to differentiate from apparent inhibition constant 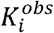. Meanwhile, *K*_*D*_ denotes similar equilibrium equation and biochemical meaning as *K*_*i*_ based on *P+ L ⇌ PL* and 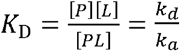, in which the dissociation constant reflects ligand concentration at the point that half of the protein is occupied under usual conditions. Consequently, *K*_*i*_ values in enzyme competitive inhibitions are convertible to the *K*_*D*_ values of proteins and small molecules if the inhibitors are structurally similar to ligands.

### 3.2 The correlation of competitive inhibition and binding affinity

Since hPNP is a well-studied enzyme with known kinetic properties and enzyme inhibition [20], hPNP and its substrate inosine and competitive inhibitor immucillin-H (ImmH) were chosen as a model of competitive enzyme inhibition. Additionally, the computational simulation on theoretical dynamic information for *K*_*m*_ and *K*_*D*_ could also be served as reference data [21-22]. Thus, it was a convenient and reliable model system for the determination of competitive enzyme inhibition.

Under strictly controlled conditions, the *K*_*m*_ value for inosine was determined to be 68.92 ± 4.08 μM, which is consistent to previous report [21]. Product formation by hPNP in the presence of ImmH exhibited a decrease of initial rates with the increase of inhibitor concentration (Fig. 1). The value of *K*_*i*_ was calculated to be 5.21 ± 1.74 nM, which is also in the accordance with previous values [22].

**Figure 1.**
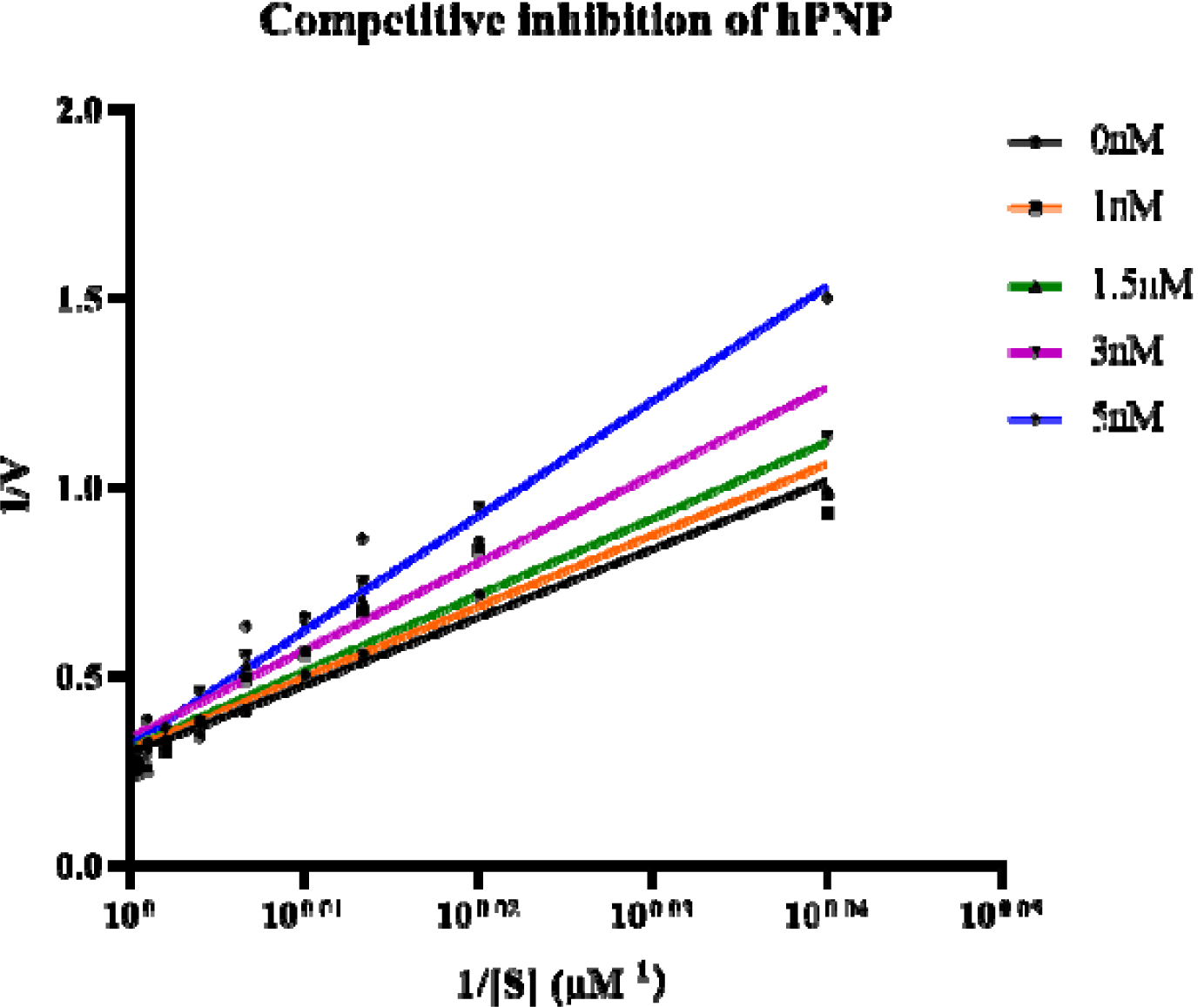
Determination of the dissociation constants for competitive inhibition of hNPN by ImmH. (5.21 ± 1.74 nM) was obtained from a replot of initial inhibition rate () as a function of inhibitor concentration.

For affinity measurements, hPNP and ImmH were used as a pair of ligand-protein in the absence of inosine (Table 1). After labeling hPNP and scanning fluorescent signal, the affinity values (*K*_*D*_) by MST were determined to be 336.40 ± 312.90 nM after fitting to MO Affinity Analysis software (Fig. 2). Next, after optimizing concentrations and titration conditions, the *K*_*D*_ values measured by ITC were 69.10 ± 18.41 nM (Fig. 3). In the case of SPR, the *K*_*D*_ values were 7.07 ± 4.23 nM from the analysis by Biacore evaluation software. (Fig. 4)

**Table 1.** The binding affinity constants of ImmH with hPNP by different assays and comparison with competitive inhibition constant^a^.

| Method | $K_d$ (nM) | $K_i$ (nM) |
| --- | --- | --- |
| | | $5.21 \pm 1.74$ |
| MST | $336.40 \pm 312.90$ | |
| ITC | $69.10 \pm 18.41$ | |
| SPR | $7.07 \pm 4.23$ | |

<sup>a</sup> Each value was determined by three independent experiments and expressed as average value with standard deviation.

**Figure 2.**
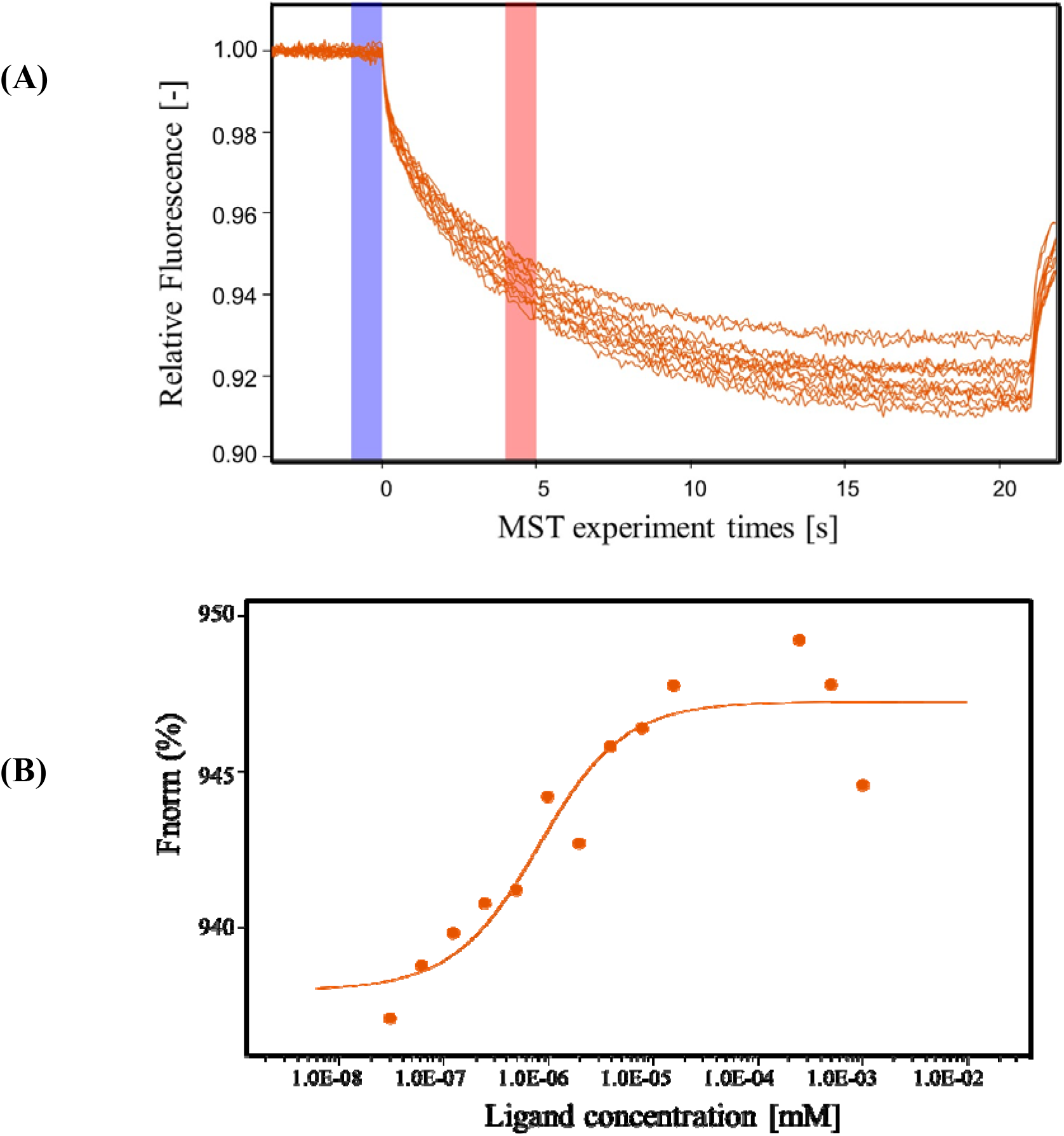
The binding affinity between hPNP and ImmH determined by MST. (A) The relative fluorescence time trace recorded by the MST instrument. (B) The binding curve of ImmH to hPNP was fitted by a standard *Kd* model.

**Figure 3.**
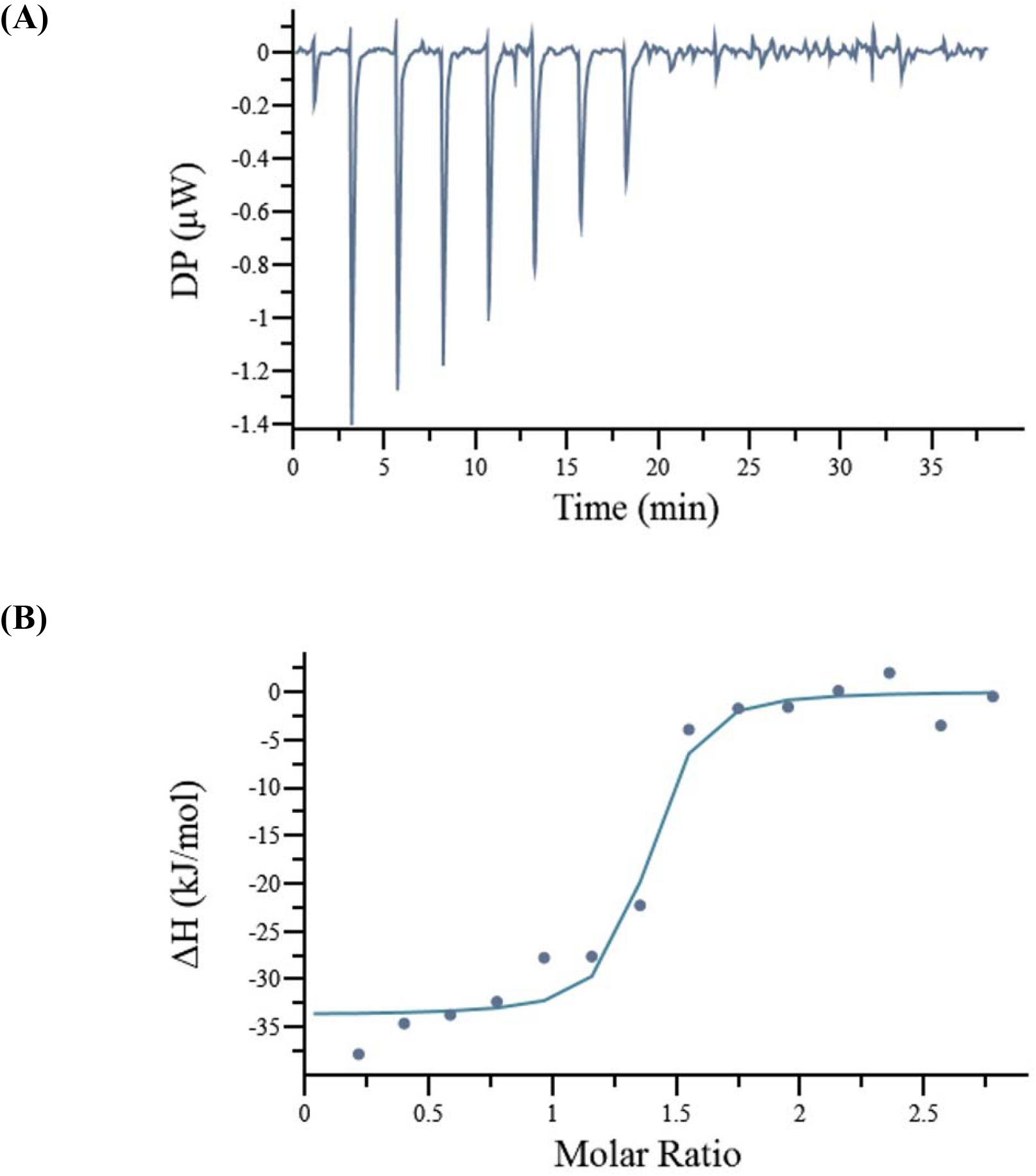
Results of the ITC measurements with ImmH and hPNP. (A) The titration of raw data after sequential injection of ImmH. (B) The integrated heat after the correlation between enthalpy changes versus the molar ratios of ImmH (150 μM) to hPNP (10 μM)) was plotted.

**Figuree 4.**
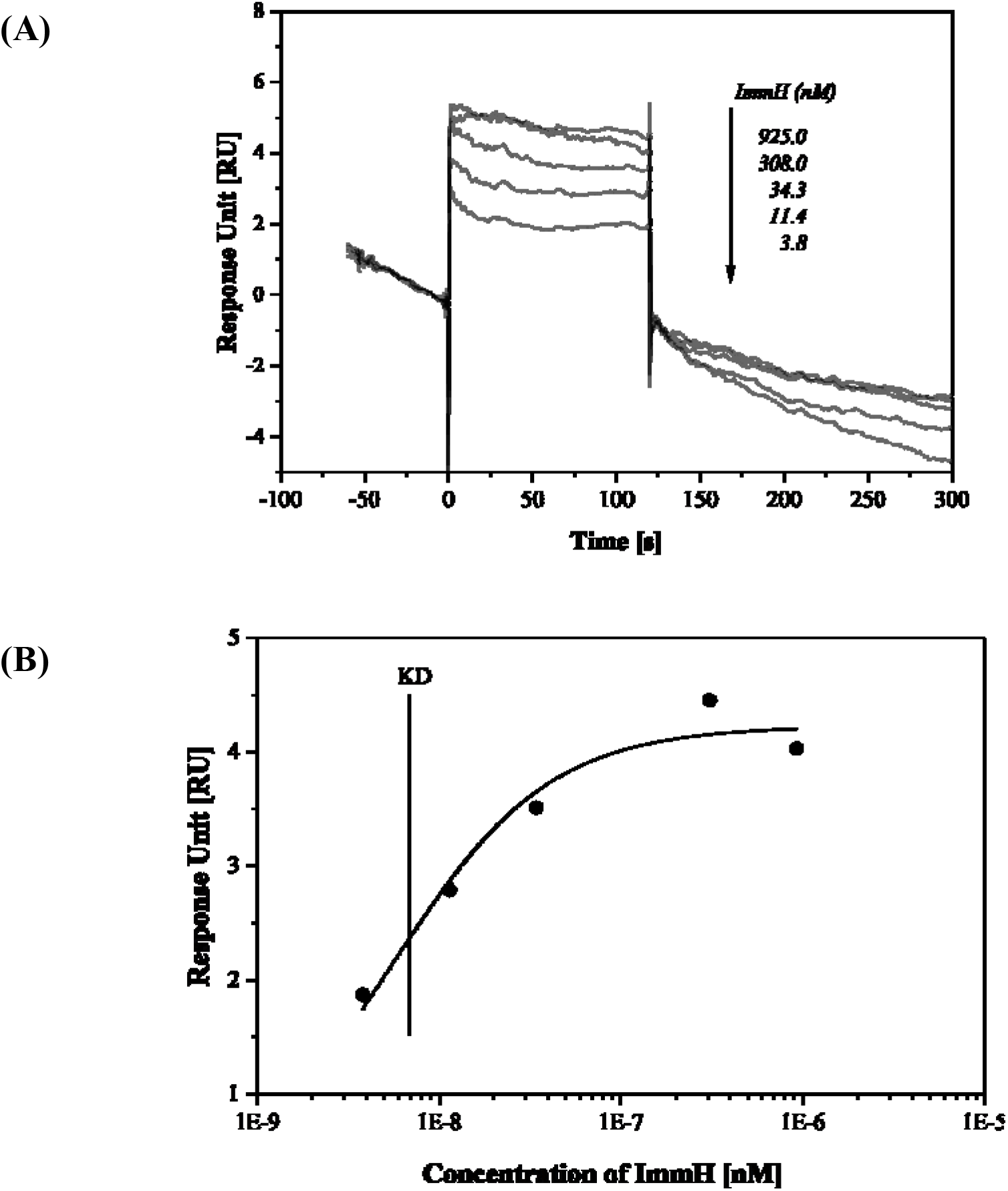
Binding affinity of ImmH to hPNP measured by SPR. (A) Plots of equilibrium response versus concentration of ImmH. (B) SPR sensorgrams by affinity analysis of the binding of ImmH to hPNP on sensor chip surface.

## 4. Discussion

According to previous studies, the determination of binding affinity between macromolecules by currently available techniques displays reasonable accuracy and reproducibility. However, in the area of small molecule inhibitor discovery and target identification, the measurements of binding affinity between a small molecule and a protein have been unreliable to result in dramatic differences by different techniques, and even a technique from different determinations. Such ambitious *K*_*D*_ data could be easily misinterpreted or misled to a wasteful effort on target identification and early drug discovery. Currently, there are three major techniques for the determination of binding affinity, namely SPR, ITC and MST, and typically only one technique would be utilized to assess *K*_*D*_ values. Therefore, there is a natural question on which technique can produce the most reliable data on biding affinity. To address this issue, a direct comparison of these techniques with a valid reference is certainly necessary. In order to have a valid reference, we speculated that competitive enzyme inhibition might be able to resemble binding affinity, and subsequently competitive *K*_*i*_ value could be used to objectively compare the *K*_*D*_ values generated by these techniques.

Although enzyme kinetics represent kinetic behaviors during the catalysis, the binding between enzyme and substrate is a prerequisite. Typically, the structure of a competitive inhibitor is very similar to that of a substrate for competing to bind the same binding site. Thus, it is highly possible to use the model of competitive enzyme inhibition to resemble the binding between a small molecule inhibitor and a protein target. Indeed, after theocratical conversion, we demonstrated the correlation between competitive *K*_*i*_ and binding affinity constant *K*_*D*_, which principally are convertible based on thermodynamic characteristics. This provided a unique opportunity for us to make a direct comparison on the accuracy and reliability of different techniques used to determine binding affinity constants. Accordingly, when the measured *K*_*D*_ value is equivalent or lower than its *K*_*i*_ value, this biding affinity constant is likely more reliable, even closer to its true affinity value. In the present study, when hPNP, inosine and ImmH were used as a model system [20-22], the *K*_*D*_ value between hPNP and ImmH was closer to the *K*_*D*_ value determined by SPR compared to MST and ITC, suggesting that real-time monitoring through resonance signal might be more suitable for detecting slight changes from the interaction of a small molecule inhibitor and a protein. The *K*_*D*_values obtained from ITC were slightly larger than *K*_*i*_ values, which could be resulted from the heat transfer effects. However, MST showed poor reproducibility on *K*_*D*_ values (approximately 64 times larger than *K*_*i*_), which could be caused by various factors [23]. SPR monitors the entire interaction process,whereas MST just compares the variation after binding. In addition, the utilization of SPR in equilibrium analysis is highly compatible with the rapid reversible binding process. Therefore, based on the complexity of the assay mixtures and processes and the differences on individual principle, the present study only provided an estimation on the accuracy of these techniques by comparative analysis regardless of the significant differences.

Given the fact that most proteins for target identification and early drug discovery are not enzymes, it would be impossible to utilize enzyme kinetics to assess binding affinity constants. Additionally, our present results indicated that currently available techniques are not able to accurately reflect the real affinity of small molecule inhibitors and proteins. Moreover, *K*_*D*_ values will continue to be an important parameter for the prediction of the draggability of small molecule inhibitors. Collectively, the present study alarms the need of developing new technique(s) for accurately, reliably and reproducibly determine the binding affinity constants for small molecule inhibitors.

## Acknowledgements

This work was supported by grants from the National Key R&D Program of China (2018YFA0902000), Key Research and Development Project of Guangdong Province (2022B1111070004), the Double “First-Class” University Project (CPU2022QZ08), and the Project Program of the State Key Laboratory of Natural Medicines, China Pharmaceutical University (SKLNMZZ202201).

---

**Scheme 1.**
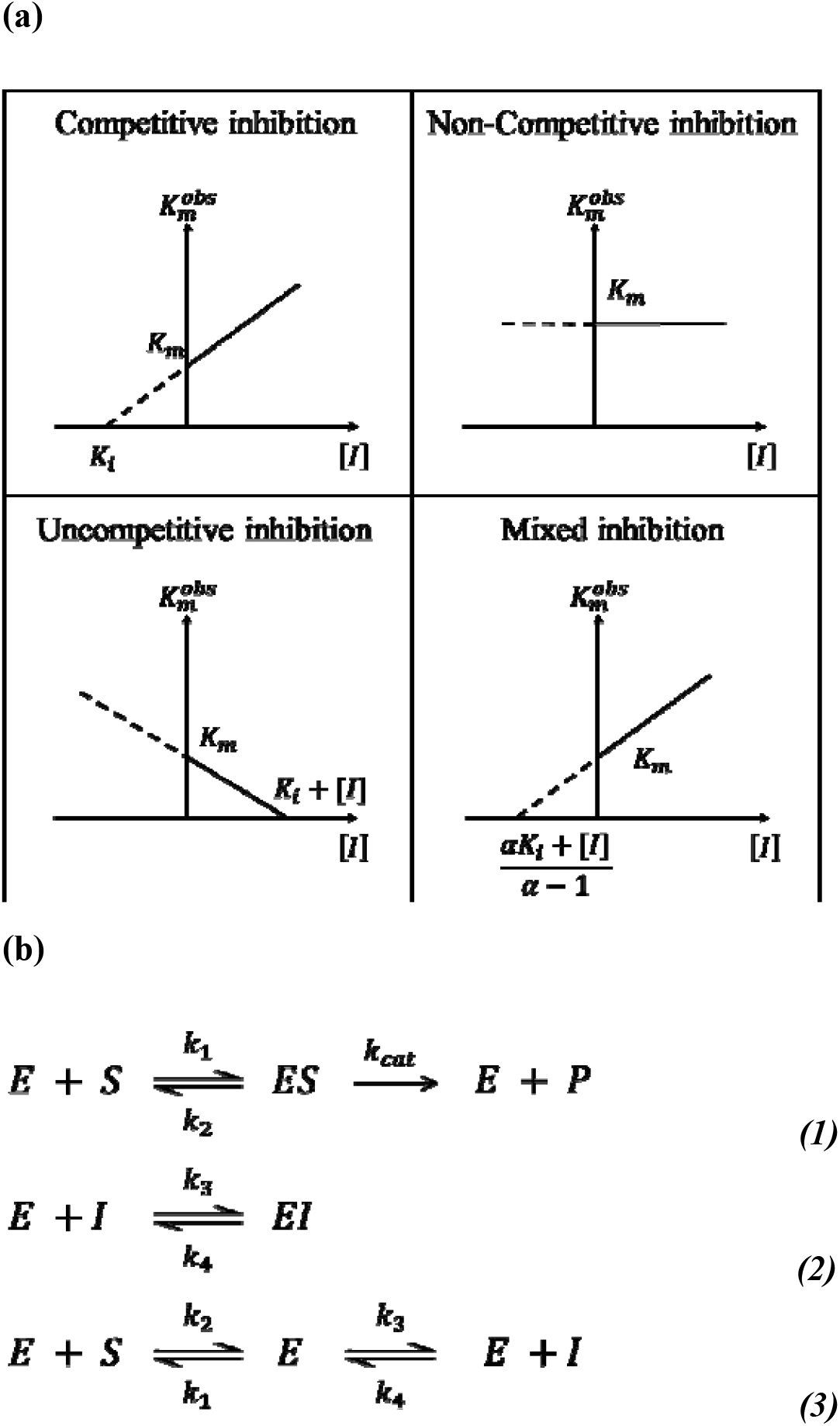
Related equilibrium equations and formulas in enzyme kinetics. (a) linear relationship between of and in different types of inhibition; (b) three equilibriums in competitive inhibition.

